# Multiple spatial behaviours govern social network positions in a wild ungulate

**DOI:** 10.1101/2020.06.04.135467

**Authors:** Gregory F Albery, Alison Morris, Sean Morris, Josephine M Pemberton, Tim H. Clutton-Brock, Daniel H Nussey, Josh A Firth

## Abstract

The structure of wild animal social systems depends on a complex combination of intrinsic and extrinsic drivers. Population structuring and spatial behaviour are key determinants of individuals’ observed social behaviour, but quantifying these spatial components alongside multiple other drivers remains difficult due to data scarcity and analytical complexity. We used a 43-year dataset detailing a wild red deer population to investigate how individuals’ spatial behaviours drive social network positioning, while simultaneously assessing other potential contributing factors. Using Integrated Nested Laplace Approximation (INLA) multi-matrix animal models, we demonstrate that social network positions are shaped by two-dimensional landscape locations, pairwise space sharing, individual range size, and spatial and temporal variation in population density, alongside smaller but detectable impacts of a selection of individual-level phenotypic traits. These results indicate strong, multifaceted spatiotemporal structuring in this society, emphasising the importance of considering multiple spatial components when investigating the causes and consequences of sociality.

**Authorship Statement:** GFA conceived the study, analysed the data, and wrote the manuscript, advised by JAF. AM and SM collected the data. JAF, JMP, THCB, and DN commented on the manuscript.

**Data Accessibility Statement:** The code used here is available at https://github.com/gfalbery/Spocial_Deer. On acceptance, the data will be uploaded to the same repo, which will be archived on Zenodo.

## Introduction

Social behaviour is an integral component of an animal’s phenotype, driving processes including disease transmission, mating, learning, and selection (Croft *et al.* 2008; VanderWaal *et al.* 2014; Krause *et al.* 2015; Firth *et al.* 2018; Sah *et al.* 2018; Silk *et al.* 2019; Firth 2020). Contemporary studies of animal behaviour often use social networks to derive individual-level social network positions, under the notion that between-individual variation in network positioning is indicative of between-individual variation in social behaviour (Franks *et al.* 2010; Krause *et al.* 2015; Sosa *et al.* 2020). However, an animal’s position in its social network is also dependent on its own spatial behaviour (Webber & Vander Wal 2018; Albery *et al.* 2020a), and on a range of extrinsic factors: demography determines local population density and structuring (Shizuka & Johnson 2019), while the environment shapes resource distributions, movement corridors, and emergent patterns of space use, all of which will influence the architecture of the social system (Firth & Sheldon 2016; Webber & Vander Wal 2018; Farine & Sheldon 2019; He *et al.* 2019). As such, it is important to consider spatial behaviour and environmental context when assessing the causes and consequences of individual-level social network positioning (Webber & Vander Wal 2018; He *et al.* 2019; Albery *et al.* 2020a), yet doing so remains difficult in most systems due to the complexity of spatial-social analyses that incorporate these processes.

The spatial drivers of social network structure are poorly understood because they are highly multivariate and (therefore) difficult to analyse. On the one hand, there is strong support for simpler “first-order” associations between spatial and social behaviour. For example, spatial proximity and social connections are often correlated, because individuals that share more space are more likely to associate or interact. This finding holds for diverse taxa including elk (Vander Wal *et al.* 2014), raccoons (Robert *et al.* 2012), birds (Firth & Sheldon 2016), and myriad other systems. Similarly, spatial and social network centrality are occasionally found to correlate (Mourier *et al.* 2019), as are temporal variation in population density and social contact rates (Sanchez & Hudgens 2015), and the social environment can drive spatial behaviour (Firth & Sheldon 2016; Spiegel *et al.* 2016). Spatial behaviours can be summarised using a wide range of metrics, including individuals’ spatial activity levels (e.g. home range area), pairwise space sharing (e.g. distances or home range overlaps), demographic structure (e.g. temporal population size or local conspecific density), and point location on the two-dimensional landscape. For example, are more social individuals simply wider-ranging, leading them to make more contacts? Do they most often inhabit areas of high population density or well-used movement corridors? These variable spatial components take a combination of different data structures, and are therefore difficult to include in the same models, particularly in large numbers and alongside a range of other individual-level phenotypes. It is therefore unclear to what extent individuals’ social network positions emerge from 1) their own social behaviour; 2) their own spatial behaviour; 3) their situation within the population and broader social network; 4) other aspects of their biotic and abiotic environment such as landscape structure; and 5) intrinsic phenotypic traits that researchers are commonly interested in investigating.

Several frameworks have been proposed to facilitate the untangling of spatial and social processes in wild animals (Jacoby & Freeman 2016; Silk *et al.* 2018, 2019; Webber & Vander Wal 2018; Mourier *et al.* 2019; Albery *et al.* 2020a). To date, statistical methodology focusses on incorporating spatial behaviours into the node-and-edge structure of network data, using e.g. null network permutations (Firth & Sheldon 2016), spatially embedded networks (Daraganova *et al.* 2012), and nested “networks of networks” composed of movement trajectories (Mourier *et al.* 2019). Many such analyses involve reducing movement patterns into some form of spatial network based on home range overlap or spatial proximity between dyads (Mourier *et al.* 2019). For example, statistical models named “animal models” can examine spatial variation by fitting such matrices as variance components, potentially alongside other dyadic similarity matrices (i.e., pairwise measures of similarity), to quantify genetic and non-genetic contributions to individuals’ phenotypes (Kruuk 2004; Stopher *et al.* 2012b; Regan *et al.* 2016; Thomson *et al.* 2018; Webber & Vander Wal 2018). As yet, the focus on controlling for spatial autocorrelation using space sharing and network permutations has contributed to a lack of clarity concerning the role that spatial behaviour and environmental context play in driving social network positioning (Albery *et al.* 2020a).

Studies across ecological disciplines increasingly use Integrated Nested Laplace Approximation (INLA) models to control for spatial autocorrelation in a multitude of contexts (Lindgren *et al.* 2011; Lindgren & Rue 2015; Zuur *et al.* 2017). As well as including fixed and random effects to quantify individual-level drivers, these models can incorporate dyadic space sharing components (Holand *et al.* 2013) and stochastic partial differentiation equation (SPDE) effects to model 2-dimensional spatial patterns in the response variable, thereby controlling for and estimating spatiotemporal variation associated with fine-scale positioning within the landscape (Albery *et al.* 2019). As such, these models offer an exciting opportunity to test and compare the roles of a range of spatial behaviours and autocorrelation structures, alongside phenotypic drivers, in determining social network positioning.

We address this question using the long-term study in the Isle of Rum red deer (*Cervus elaphus*). These study animals comprise an unmanaged wild population with a contiguous fission-fusion social system (Clutton-Brock *et al.* 1982). They experience strong environmental gradients and exhibit spatial autocorrelation in a number of important phenotypes: individuals with greater home range overlap have more similar behavioural and life history traits (Stopher *et al.* 2012b), and those in closer proximity have more similar parasite burdens (Albery *et al.* 2019); further, as with other matrilineal mammalian systems, closely related individuals frequently associate (Clutton-Brock *et al.* 1982) and live closer together (Stopher *et al.* 2012b). Individuals have highly repeatable home ranges (Stopher *et al.* 2012b) that decline in size over their lifetimes, predicting declining survival probability (Froy *et al.* 2018). The study area has a strong spatial gradient in resource availability, with high-quality grazing heavily concentrated in the far north of the system, and with most individuals aggregating around this area, such that population density decreases outwards towards the edge of the study population (Clutton-Brock *et al.* 1982). As such, the deer comprise an ideal system for assessing spatial-social relationships in the wild.

To assess how individuals’ spatial behaviours translate to social network positions, we constructed fine-scale social networks from 43 years of censuses of the study population. We derived 8 different individual-level network positioning measures of varying complexity that are important to different social processes (Krause *et al.* 2015; Sosa *et al.* 2020). Using multi-matrix animal models in INLA, we examined whether spatial locations, space sharing, home range area, and local population density explained variation in network position metrics, alongside a range of individual-, temporal-, and population-level factors. Specifically, we aimed to test two hypotheses: that the structure of the social network would be highly dependent on the distribution of population density in space; and that individuals’ social network centrality would be largely explained by their ranging behaviour, where wide-ranging individuals were more likely to be socially well-connected. We further expected that space sharing and point locations would uncover substantial spatial autocorrelation in social network positioning, and that different social network metrics would exhibit different spatial patterns and vary drivers. This not only comprises a large-scale empirical examination of the factors shaping social network positions in this extensively monitored wild mammal, but also provides a methodological advancement in developing powerful, flexible new methods (INLA-based multi-matrix animal models) with broad potential for examining spatial-social processes in this and other systems.

## Methods

### Study system and censusing

The study was carried out on an unpredated long-term study population of red deer on the Isle of Rum, Scotland (57°N,6°20’W). The natural history of this matrilineal mammalian system has been studied extensively (Clutton-Brock et al. 1982), and we focussed on females aged 3+ years, as these individuals have the most complete associated census data, and few males live in the study area except during the mating period. Individuals are monitored from birth, providing substantial life history and behavioural data, and >90% of calves are caught and tagged, with tissue samples taken (Clutton-Brock *et al.* 1982). The population thus has comprehensive genomic data, allowing high-powered quantitative genetic analyses: most individuals born since 1982 have been genotyped at >37,000 SNPs, distributed throughout the genome (e.g. Huisman, Kruuk, Ellis, Clutton-Brock, & Pemberton, 2016). Census data were collected for the years 1974-2017, totalling 423,070 census observations. Deer were censused by field workers five times a month, for eight months of the year, along one of two alternating routes (Clutton-Brock *et al.* 1982). Individuals’ identities, locations (to the nearest 100M), and group membership were recorded. Grouping events were estimated by seasoned field workers according to a variant of the “chain rule” (e.g. Castles et al., 2014), where individuals grazing in a contiguous group within close proximity of each other (under ~10 metres) were deemed to be associating, with mean 130.4 groups observed per individual across their lifetime (range 6-943). The mortality period falls between Jan-March, when there is least available food, and minimal mortality occurs outside this period. We only used census records in each May-December period, from which we derived annual social network position measures as response variables (Figure 1-2). We elected to investigate this seasonal period because it stretches from the spring calving period until the beginning of the mortality period, simplifying network construction and avoiding complications arising from mortality events. Our dataset totalled 3356 annual observations among 532 grown females (Figure 1).

**Figure 1:**
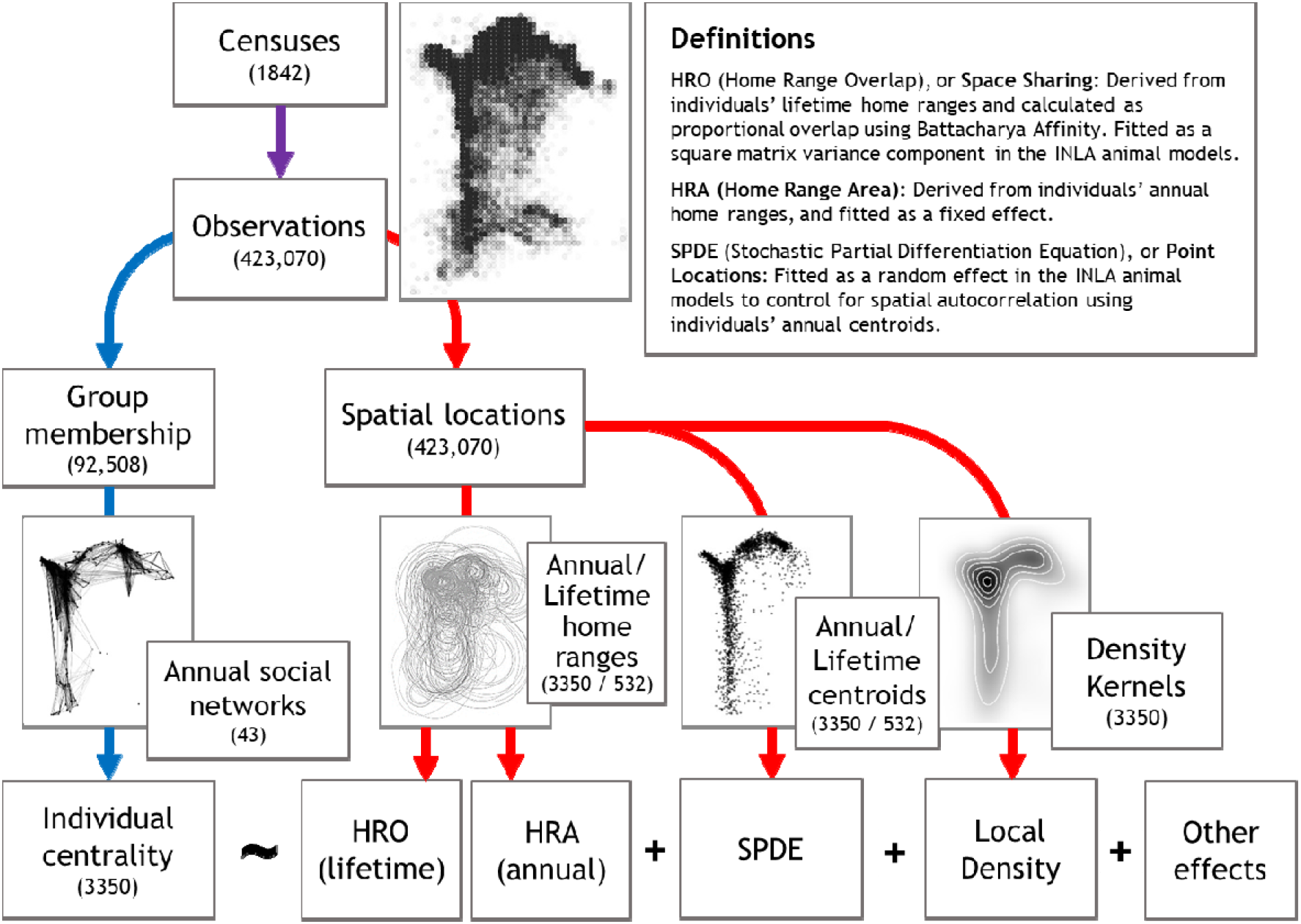
Data processing and analysis pipeline, demonstrating how behavioural census data were collected, used to derive social and spatial behavioural traits, and fitted in INLA animal model GLMMs. Numbers in brackets represent sample sizes, and only include females aged 3+ years. Blue arrows represent social behaviour; red arrows represent spatial behaviours. See methods for the fixed and random effects. The text box displays the definitions for the different spatial effects.

**Figure 2:**
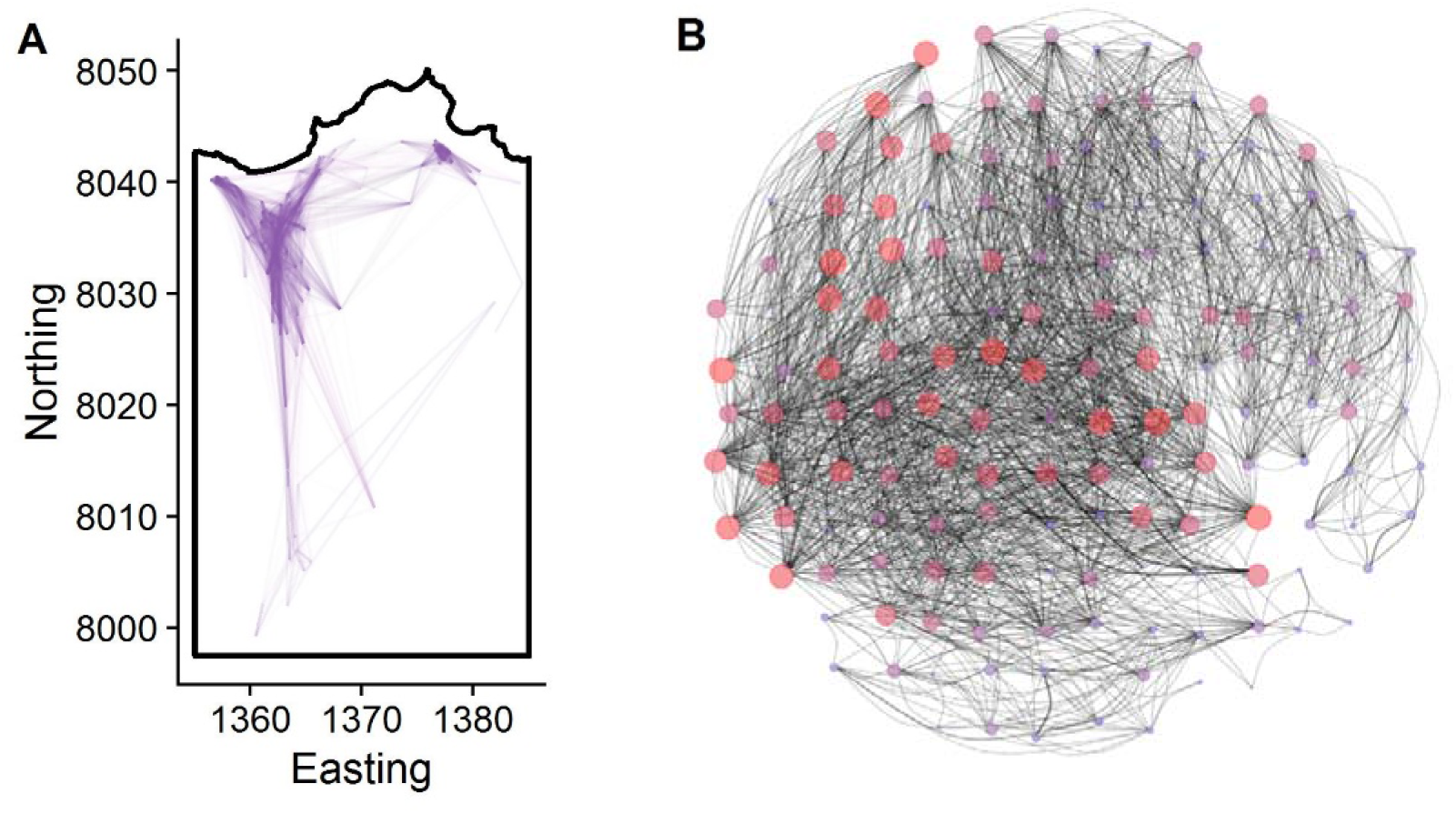
Spatial structuring of the 2016 social association network as a representative example. A: the spatial locations (centroids) of individual deer, connected by their social associations. Line opacity and width are weighted by connection strength. Ten axis units = 1KM. B: the same social network with the nodes positioned in a network spring-layout (Csardi & Nepusz 2006) and then expanded into an even, circular grid according to their nearest spatial positions in A. The points’ (i.e. nodes’) sizes and colours show individuals’ strength centrality (large and red=high strength; small and blue=low strength). Thickness of the lines (i.e. edges) connecting them shows dyadic association strength between individuals.

In this system, female reproduction imposes substantial costs for immunity and parasitism (Albery *et al.* 2020c), and for subsequent survival and reproduction (Clutton-Brock, Albon, & Guinness, 1989; Froy, Walling, Pemberton, Clutton-Brock, & Kruuk, 2016). If a female reproduces, she produces 1 calf per year in the spring, generally beginning in May; the “deer year” begins on May 1 for this reason. Here, reproductive status was classified into the following four categories using behavioural observations: True Yeld (did not give birth); Summer Yeld (the female’s calf died in the summer, before 1^st^ October); Winter Yeld (the female’s calf died in the winter, after 1^st^ October); and Milk (calf survived to 1^st^ May the following calendar year).

### Generating spatial and social matrices

All code is available online at https://github.com/gfalbery/Spocial_Deer. We constructed the home range overlap (HRO) matrix using the R package AdeHabitatHR (Calenge 2011), following previous methodology (Stopher *et al.* 2012b; Regan *et al.* 2016; Froy *et al.* 2018).

First, using a kernel density estimation method, we derived lifetime home ranges for each individual with more than five census observations. Previous analysis has shown that this system is robust to the number of observations used to generate home ranges (Froy *et al.* 2018). We used lifetime home ranges to fit one value per individual in the animal models; individual ranges (and range sizes) correlate strongly from year to year (Stopher *et al.* 2012b; Froy *et al.* 2018). We derived proportional HRO of each dyad using Bhattacharya Affinity (following Stopher *et al.* 2012b), producing values between 0-1 (i.e. no overlap to complete overlap).

To control for individuals’ two-dimensional point locations, we used a Stochastic Partial Differentiation Equation (SPDE) effect in INLA. This effect models the distance between points to calculate spatial autocorrelation, using Matern covariance (Lindgren *et al.* 2011). This random effect used individuals’ annual centroids (mean easting and northing in a given year) or lifetime centroids (mean easting and northing across all observations) as point locations to approximate spatial variation in the response variable (Lindgren *et al.* 2011; Albery *et al.* 2019).

We used a genomic relatedness matrix (GRM) using homozygosity at 37,000 Single Nucleotide Polymorphisms, scaled at the population level (Yang *et al.* 2011; for a population-specific summary, see Huisman *et al.* 2016). This matrix is well-correlated with pedigree-derived relatedness metrics (Huisman *et al.* 2016). HRO was well-correlated with distance between lifetime centroids (i.e., closer individuals tended to share more range), and both were weakly but significantly correlated with genetic relatedness (Supplementary Figure 1).

To test whether social network positions could be explained by population density, we derived the local density of individuals again using AdeHabitatHR (Calenge 2011). We generated density kernels of observations, and then assigned individual deer their local population density based on their location on this kernel, following previous methodology developed in badgers (Albery *et al.* 2020b). This local density value was then fitted as a fixed explanatory variable. We used four different density metrics, each examining the density of a different observation type: lifetime centroids (“lifetime density”); annual centroids (“annual density”); all observations across the study period (“sighting density”); and all observations in the focal year (“annual sighting density”). Only one such density metric was fitted at once. We also calculated annual home range areas (HRA) by taking the 70% isopleth of each individual’s annual space use distribution, following previous methodology (Froy *et al.* 2018). This HRA variable was fitted as a fixed effect in the same way as local density.

We constructed a series of 43 annual social networks using “gambit of the group,” where individuals in the same grouping event (as described above) were taken to be associating (Franks *et al.* 2010). Dyadic associations were calculated using the ‘simple ratio index’ (Cairns & Schwager 1987) derived as a proportion of total sightings (grouping events) in which the focal individuals were seen together: Sightings_A,B_/(Sightings_A_+Sightings_B_-Sightings_A,B_), or Intersect_A,B_/Union_A,B_. In this dyadic matrix, 0=never seen together and 1=never seen apart.

### Statistical Analysis

#### Metrics

Using the annual social networks, we derived eight individual-level network metrics which are commonly used across animal social networks and have been considered in detail(Whitehead 2008; Brent 2015; Krause *et al.* 2015; Firth *et al.* 2017). We set each of these network metrics for use as response variables in separate INLA Generalised Linear Mixed Models (GLMMs) with a Gaussian family specification. In increasing order of complexity, our measures included four direct sociality metrics, which only take into account an individual’s connections with other individuals: 1) Group Size – the average number of individuals a deer associated with per sighting; 2) Degree – the number of unique individuals she was observed with; 3) Strength – sum of all their weighted social associations to others; 4) Mean Strength – the average association strength to each of the unique individuals she was observed with (equivalent to strength divided by degree). We also included four more complex “indirect” metrics (all using algorithms as specified from (Csardi & Nepusz 2006)), which also take into account an individual’s connections’ connections: 5) Eigenvector centrality – which considers the sum of their own connections and the sum of their associates’ connections; 6) Weighted Eigenvector – which is akin to eigenvector centrality but also accounts for the weights of theirs, and their associates, connections; 7) Betweenness – the number of shortest paths that pass through the focal individual to traverse the whole network; 8) Clustering (local) – the tendency for an individual’s contacts to be connected to one another, forming triads. The raw, untransformed correlations were assessed for all metrics, and R lay between −0.5 and 0.879 across metrics (Supplementary Figure 2). When modelling them as response variables, to approximate normality, all social metrics were square root-transformed apart from eigenvector centralities (which were left untransformed), group size (which was cube root-transformed), and betweenness (which was log(X+1)-transformed). Each social network metric was fitted as a response variable in a separate model set (as outlined conceptually in Figure 1).

#### Base model structure

We ensured that all models followed the same base structure. Random effects included individual identity and year (categorical random intercepts), as well as the genetic relatedness matrix. Fixed effects included Age (continuous, in years), Reproductive Status (four categories: True Yeld; Summer Yeld; Winter Yeld; and Milk), and Number of observations (continuous, log-transformed to approximate normality), as well as year-level continuous factors including Year (continuous) and that year’s study Population Size (log-transformed). All continuous response and explanatory variables were standardised to have a mean of zero and a standard deviation of 1. Fixed effect estimates were provided by the mean and 95% credibility intervals of the posterior estimate distribution.

#### Adding spatial components

To investigate the divergent value of different spatial behaviours, we iteratively added spatial effects to the base model, investigating which behaviours best fit the data. These spatial behaviours corresponded to four broad components in Figure 1: space sharing (HRO matrix); home range area (HRA); point locations (SPDE effect); and local population density (density fixed effect). For space sharing, we only used one metric: lifetime HRO (see above). For point locations, we selected between 1) lifetime centroids; 2) annual centroids; and 3) annual centroids with a spatiotemporally varying annual field. For density, we used the four metrics outlined above (“lifetime”, “annual”, “sighting”, and “annual sighting” density). To distinguish between competitive models we used Deviance Information Criterion (DIC). In each round, we added each spatial behaviour individually and then kept the best-fitting one, until all four had been added or their addition did not improve the model, using a cutoff of 2 DIC.

#### Comparing all spatial and non-spatial drivers

To compare the relative importance of all fixed and random effects, we examined the model’s predicted values and their correlations with the observed values, representing the proportion of the variance that was explained by the model (i.e., R^2^). We used the model to predict each social behaviour metric, and iteratively held each explanatory variable’s predictions at the mean, one at a time. We then assessed the squared correlations of these values with the observed values, relative to those of the full model. Variables with greater effects in the model produced less accurate predicted values when held constant.

## Results

Spatial behaviours were important in determining all eight individual-level social network position variables. The non-spatial model was far the poorest-fitting for all eight metrics, and the DIC changes associated with adding spatial components were substantial (Figure 3A). Generally, wide-ranging individuals and those living in areas of greater population density tended to be more central, and space sharing and point location effects both revealed substantial spatial autocorrelation (Figure 3).

**Figure 3:**
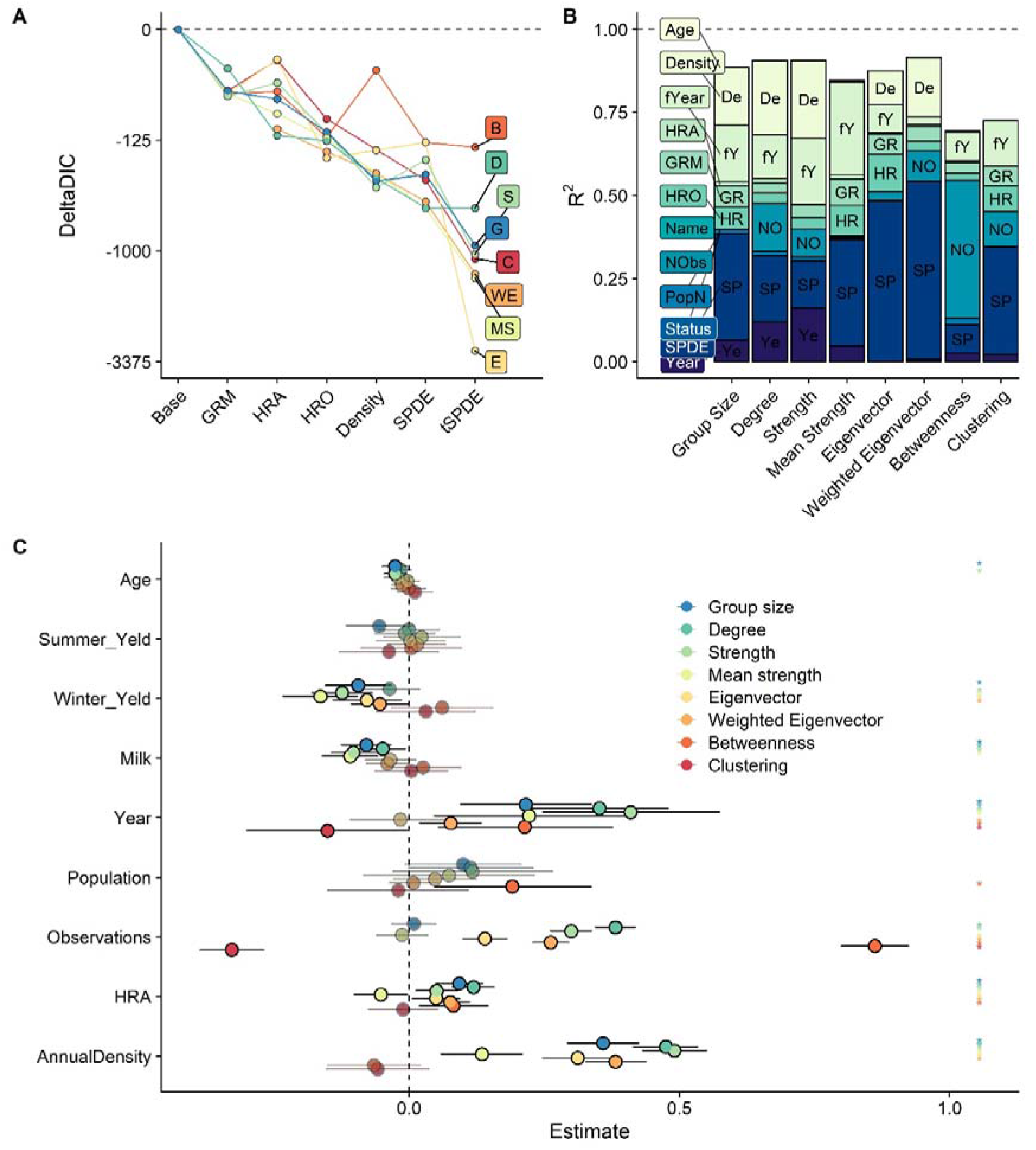
Model outputs demonstrating strong effects of spatial and non-spatial drivers on social network positions. A: DIC changes associated with addition of different spatial components, for all eight social network centrality measures. Variables are arranged in order of mean contribution to model fit, which varied little among response variables. Different colours correspond to different network centrality response variables, with the same colour key as panel C. GRM = Genomic Relatedness Matrix. HRA = Home Range Area. HRO = home range overlap. The SPDE models are differentiated into those using annual centroids (“SPDE”) and the version with spatiotemporally varying annual spatial fields (“tSPDE”). B: Variance accounted for by each variable for all eight network position measures, expressed as contribution to R^2^ in the annual model (squared correlation between observed and predicted values). Different shades correspond to different variables. fYear = year as a categorical random effect. HRA = Home Range Area. GRM = Genomic Relatedness Matrix. HRO = home range overlap. Name = individual identity. NObs = number of observations (i.e., sampling bias). PopN = population size. Status = reproductive status. SPDE = point location effects estimated using the Stochastic Partial Differentiation Equation effect in the INLA models. For all response variables, individual level effects (Age, Reproductive Status, Name) had a negligible effect. C: Fixed effect estimates for the models. Fixed effects are grouped into individual factors (age and three reproductive status effects), annual factors (continuous time in years since study began, and annual population size), and sampling factors (observation number). Reproductive status effects are separated into four levels: did not reproduce (the intercept); calf died in the first few months of life (“Summer Yeld”); calf died during the winter (“Winter Yeld”); and calf survived to May the following year (“Milk”). Different colours correspond to different network centrality response variables. Points represent the posterior mean; error bars denote the 95% credibility intervals for the effects. Asterisks denote significant variables (i.e., those whose estimates did not overlap with zero). Significant variables are fully opaque, while non-significant ones are transparent.

As expected, home range area and population density had substantial effects on social network centrality (Figure 3). Population density was positively associated with all centrality measures except betweenness and clustering (Figure 3A), and the best-fitting density metric was annual density. Individuals with larger home ranges likewise tended to be more social, except in the case of clustering (no effect) and mean strength, which were negatively associated with HRA (Figure 3A).

Notably, point location-based SPDE effects tended to improve model fit over these fixed effects, and had a greater effect than on model fit space sharing HRO effects, even when conceptualised at the same timescale (i.e., across the individual’s lifetime). Investigating the R^2^ components of the models containing only HRO (i.e., without SPDE effects) revealed that in general spatial overlap accounted for more variation than the genetic matrix (Supplementary Figure 3), but comparing these with the other models revealed that the point location effects contributed more than either of these matrices (Figure 3B). Annually varying centroids further improved model fit, and allowing the spatial field to vary between years in our spatiotemporal models improved models even more (Figure 3A). Although the space sharing and genomic relatedness matrices had similar sized impacts on the full models (Figure 3B), removing the SPDE effect resulted in a substantial increase in the HRO effect, but with very little impact on the GRM’s R^2^ (Supplementary Figure 3). These findings were relatively consistent across all metrics (Figure 3A-B), although the SPDE effect was notably smaller for betweenness (Figure 3B). Taken together, these results reveal that lifetime space sharing was good at accounting for variation in social behaviour, but that its effect was surpassed by increasingly complex temporal formulations of point location effects.

We compared the importance of all fixed and random effects by predicting selectively from the model, revealing overwhelmingly strong effects of spatiotemporal drivers (Figure 3B). Our models fit well and explained a substantial amount of variation in social network centrality (>70%); the majority of the fit was lent by a combination of the INLA SPDE effect, fixed effects of local population density, and random effects of year (Figure 3B). Space sharing (HRO) and home range area (HRA) had comparatively small effects.

Observations also had a notable impact for Degree, Betweenness, and Clustering (Figure 3B). Fixed effects for year and observation numbers were generally strong and significantly positive across metrics, except in the case of clustering, for which observation number’s effect was significantly negative (Figure 3B). There were also small positive effects of population size on betweenness and degree centrality (Figure 3B).

Although individual-level drivers (reproduction, age, and individual identity) had a negligible impact on all variables’ R^2^ (Figure 3B), many had a significant effect (i.e., their 95% credibility intervals did not overlap with zero; Figure 3C). Individuals whose calves lived to the winter and then either died before the 1^st^ May (“Winter Yeld”) or survived (“Milk”) were generally less central than those that did not give birth (“True Yeld”) or whose calf died before 1^st^ October (“Summer Yeld”). Similarly, there were minor age-related decreases in network centrality for the direct metrics (Group Size, Degree, and Strength; Figure 3C).

To investigate spatial patterns of sociality when accounting for our fixed and random effects, we projected the annual SPDE random effect in two-dimensional space (Figure 4; Supplementary Figures 5-12). The spatial distributions of network centrality metrics were highly variable, but direct metrics generally peaked in the centre of the study system and decreased outwards (Figure 4). Mean Strength was an exception, being lowest in the centre and increasing outward (Figure 4D); Clustering was patchily distributed, such that no clear pattern was evident (Figure 4H); and Betweenness was slightly offset, being highest in the north-northeast of the study area rather than in the central north (Figure 4G). The range of autocorrelation also varied among metrics; Betweenness and Clustering had notably shorter ranges than the other metrics (Supplementary Figure 4). We also plotted the spatial fields through time, revealing substantial variation in the spatial fields across the study period (Supplementary Figures 5-12).

**Figure 4:**
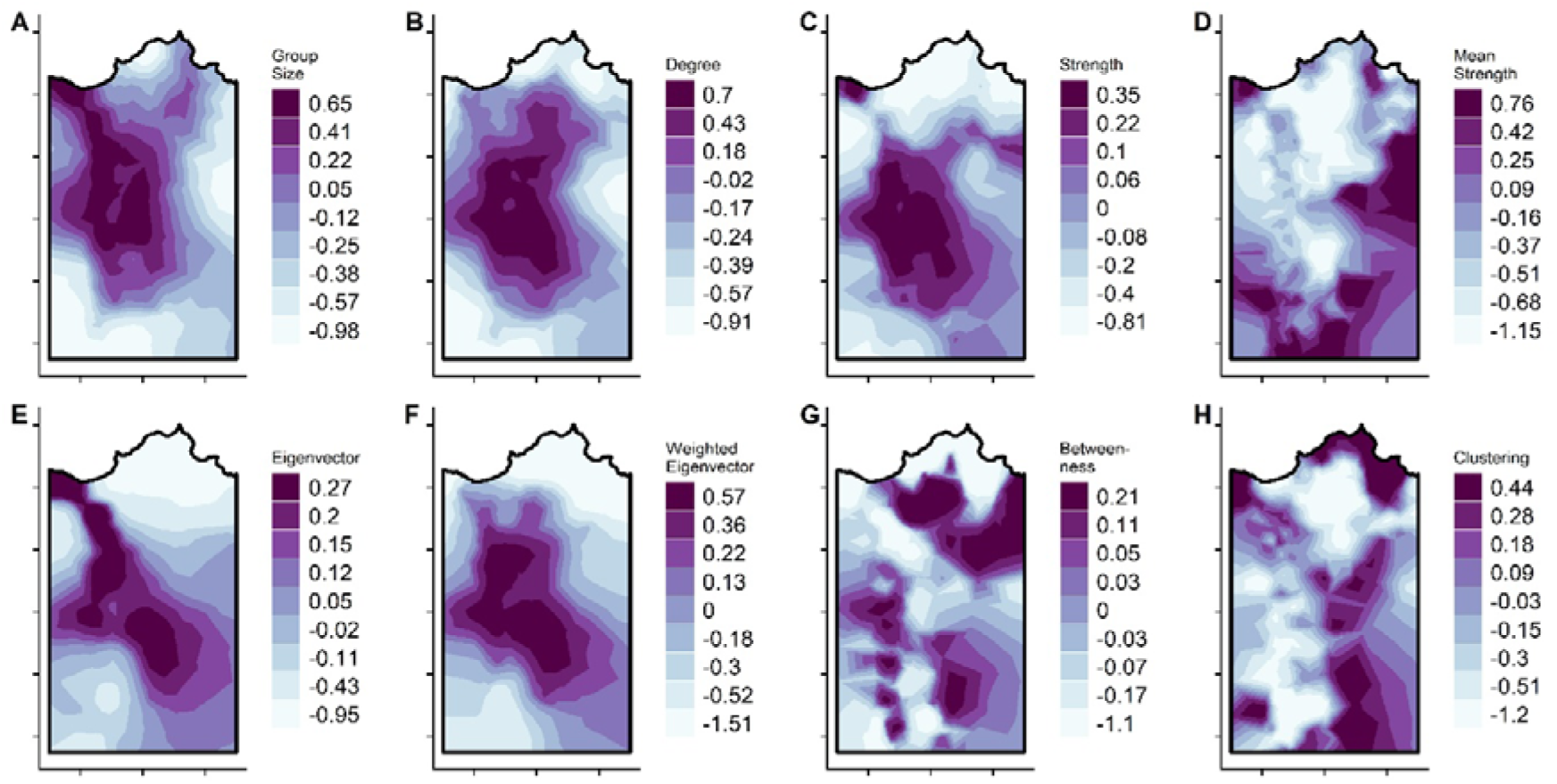
Spatial fields for the SPDE random effect for each response variable, taken from the INLA animal models and based on annual centroid point locations. Metrics can be conceptualised as simpler “direct” metrics (top row) and more complex “indirect” metrics (bottom row). Darker colours correspond to greater values. Each axis tick corresponds to 1km; for the values associated with the Easting and Northings, see Figure 1.

## Discussion

### The role of spatial behaviour in driving social network structure

The position individuals occupy within their social networks can affect many aspects of their ecology and evolution (Krause *et al.* 2015; Firth *et al.* 2018; Sah *et al.* 2018), and our results confirm the powerful role of fine-scale spatial context in shaping such traits (e.g. Farine & Sheldon, 2019; Mourier et al., 2019; Webber & Vander Wal, 2018). Capitalising on our models’ ability to compare the influence of a wide range of spatial and non-spatial components, we found that spatial behaviour and environmental context were the most important determinants of social network centrality -- more so than a suite of individual-level phenotypes and demographic factors. Individuals with larger ranges and inhabiting higher-density areas were more central in the social network, revealing the important role of individual spatial activity levels and location within the broader population structure. As expected, models were further improved when we incorporated pairwise space sharing and two-dimensional point locations, demonstrating that an individual’s social network position is not determined simply by the density of nearby individuals and by its own spatial activity, but by other aspects of the fine-scale surrounding environment such as microclimate, resource distribution, and landscape architecture (Spiegel *et al.* 2018; Webber & Vander Wal 2018; He *et al.* 2019). Reciprocally, individuals may be altering their spatial behaviour, e.g. opting to share more space or live closer together if they are more socially connected (Firth & Sheldon 2016; Spiegel *et al.* 2016). As such, we propose that social network studies should more regularly incorporate both space sharing and (temporally varying) point locations in their statistical approaches to anticipate these effects, alongside specific spatial behaviours thought to drive social network position. This practice will help to buffer for the fact that the spatial environment not only correlates with social proximity, but can alter the fabric of the network itself.

### The landscape of sociality

One of the foremost advantages of our approach is the ability to flexibly investigate two-dimensional spatial patterns of social network centrality. This allowed us to qualitatively assess the spatial structure of the social network, while providing clues towards the causal factors. Most notably, betweenness peaked in the north-northeast of the system, likely because the far northeastern community is relatively isolated from the rest of the population due to the landscape structure (Figure 2), so that many ‘social paths’ that traverse the population (the criteria for betweenness centrality) go through individuals in this intermediate (north-northeast) area. That is, individuals living in this area are more likely to be connected to both the far eastern communities and the central and western ones.

As expected, direct centrality metrics (group size, degree, and strength) were affected by local population density, which peaks in the central north study area due to the concentration of high quality grazing (Clutton-Brock *et al.* 1982). Individuals’ resource selection behaviours increase local density in this area (Clutton-Brock *et al.* 1982), and will increase social connectivity as a result (Ostfeld *et al.* 1986; Sanchez & Hudgens 2015; Webber & Vander Wal 2018). This comprises strong evidence for density-related increases in social contact frequency, and accentuates the vital importance of considering resource distribution, habitat selection, and population structure when examining social network correlates (Spiegel *et al.* 2016; Webber & Vander Wal 2018; Farine & Sheldon 2019; He *et al.* 2019). However, because density was accounted for as a fixed effect in the models, the spatial patterns of location effects for the direct metrics did not strictly follow the spatial pattern of density. Instead, these metrics peaked in the centre of the study population, demonstrating that individuals living in this central region are more well-connected *when accounting for population density*. Combining these spatial components allowed us to effectively differentiate what we do know (that greater population density drives increased social connectedness) from what we do not (the drivers of greater sociality for individuals in the central area). Without using the SPDE effect (i.e., relying only on generalised pairwise space sharing rather than accounting for specific two-dimensional spatial patterns), these insights into these patterns may have been harder to detect. An alternative method could involve splitting the population into subpopulations and analysing them separately or comparing them, but this method has been shown to be less powerful in this population (Albery *et al*. 2019), and is ultimately based on arbitrary choices if a population is mixed. The causes of the spatial distribution of clustering remain unresolved, but the pattern highlights areas where individuals are connected together in triads or tight cliques, and appears to be negatively correlated with betweenness (Figure 4). For traits such as this, it is unlikely that a simpler explanatory variable could be formulated to quantify the spatial-social processes at play.

Regardless of the causes of the spatial patterns, such fine-scale variation across the landscape holds important ecological consequences, particularly for the more complex network metrics. For instance, the areas of high clustering may act as ‘incubator’ areas where cliques can develop new socially influenced behaviours (Centola 2018; Guilbeault *et al.* 2018; Firth 2020) such as cooperative behaviours (Rand *et al.* 2011). The high contact rates in the northern central areas might sustain high local burdens of directly transmitted diseases (Cote & Poulin 1995), while individuals inhabiting the high-betweenness intermediate areas may be important for transmitting novel diseases across the population as a whole (VanderWaal *et al.* 2014).

### Analytical benefits of INLA animal models

Analyses using multiple layers of different behaviours are well-suited to extricating space and sociality in wild animal systems (Silk *et al.* 2018; Webber & Vander Wal 2018; Finn *et al.* 2019), and there is increasing conceptual and analytical overlap with the related field of movement ecology (Jacoby & Freeman 2016; Mourier *et al.* 2019; Pasquaretta *et al.* 2020). Notably, many spatial-social studies suffer from the necessity to reduce complex movement patterns into simpler metrics, which risks losing important information in the process. As such, recent studies have pushed for researchers to incorporate movement trajectories themselves into complex network data structures (Mourier *et al.* 2019). Our approach allows incorporation of multiple dyadic and non-dyadic behavioural measures, and with several analytical timescales, offering an alternative workaround to this problem. Although other methods can control for point locations (e.g. using autoregressive processes and row/column effects; Stopher *et al.* 2012b), INLA models allow greater precision, fit quickly, and allow incorporation of spatiotemporal structuring. Furthermore, plotting the SPDE effect in two dimensions, as in Figure 4, gives an easily interpretable and intuitive portrayal of network traits in space that can be hard to visualise using other methods. For these reasons, we highly recommend further exploration of INLA animal models as a flexible method with which to extricate individual, demographic, spatial, and temporal contributors to sociality where sample sizes are sufficient (Thomson *et al.* 2018; Webber & Vander Wal 2018). In addition to carrying out network-level manipulations (Daraganova *et al.* 2012; Davis *et al.* 2015; Firth & Sheldon 2016; Farine 2017), researchers concerned about spatial confounding could implement relatively familiar linear models of social behaviour, but with additional spatial components such as SPDE random effects and similarity matrix variance components, with trustworthy and interpretable results (Albery *et al.* 2020a).

Although space accounted for an overwhelming amount of variation, many non-spatial factors had substantial effects. The categorical random effect for interannual variation was substantial, and there were detectable linear annual effects and population size effects, as expected given the important roles of demography in structuring social networks (Shizuka & Johnson 2019). Interestingly, there was a substantial positive association with study year that was not attributable to the growth in population size over the same period. It is possible that this represents a change in the deer’s social phenotypes over time, although the potential specific mechanisms now would benefit from further examination. Individual-level factors had weaker contributions to model fit and smaller effect sizes: most notably, genetic and individual random effects were negligible when spatial autocorrelation was accounted for, confirming the importance of considering space when assessing heritability independently of space in this population (Stopher *et al.* 2012a). Nevertheless, individual-level effects were encouragingly still detectable and significant, particularly for simpler “direct” metrics. It is possible that more complex social network positions are less determined by individual social behaviours, particularly for animals with fission-fusion societies such as the deer; this hypothesis could be tested using similar spatial-social analyses in a number of other systems. This finding demonstrates that even when spatial structuring plays a vital role in determining social network structure, controlling for this structuring analytically can reveal important, conservative individual-level effects. Future analyses within this population, and potentially other long-term studies, could take advantage of this framework by including environmental drivers such as food availability and climatic factors to explain patterns of social connectivity, while further unpicking the causes of the individual-level trends that we observed.

## Acknowledgements

We thank Scottish Natural Heritage and its predecessors for permission to work on the Isle of Rum NNR. The field project has been supported by grants mainly from the UK NERC with some additional funding from BBSRC, the Royal Society and ERC. We thank all who have contributed to the maintenance of the project over time, especially Loeske Kruuk. We thank multiple dedicated field workers who have contributed to field data collection, especially Fiona Guinness who collected the first 20 years of census data. GFA was funded by NSF grant number 1414296, and by a Bruce McEwen Career Development Fellowship the Animal Models for the Social Dimensions of Health and Aging Research Network (NIH/NIH R24 AG065172). JAF was supported by a fellowship from Merton College and BBSRC (BB/S009752/1) and funding from NERC (NE/S010335/1). We thank Amy Sweeny and Quinn Webber for comments on the manuscript, as well as Matt Silk, Orr Spiegel, and one anonymous reviewer.

---

**Supplementary Figure 1:**
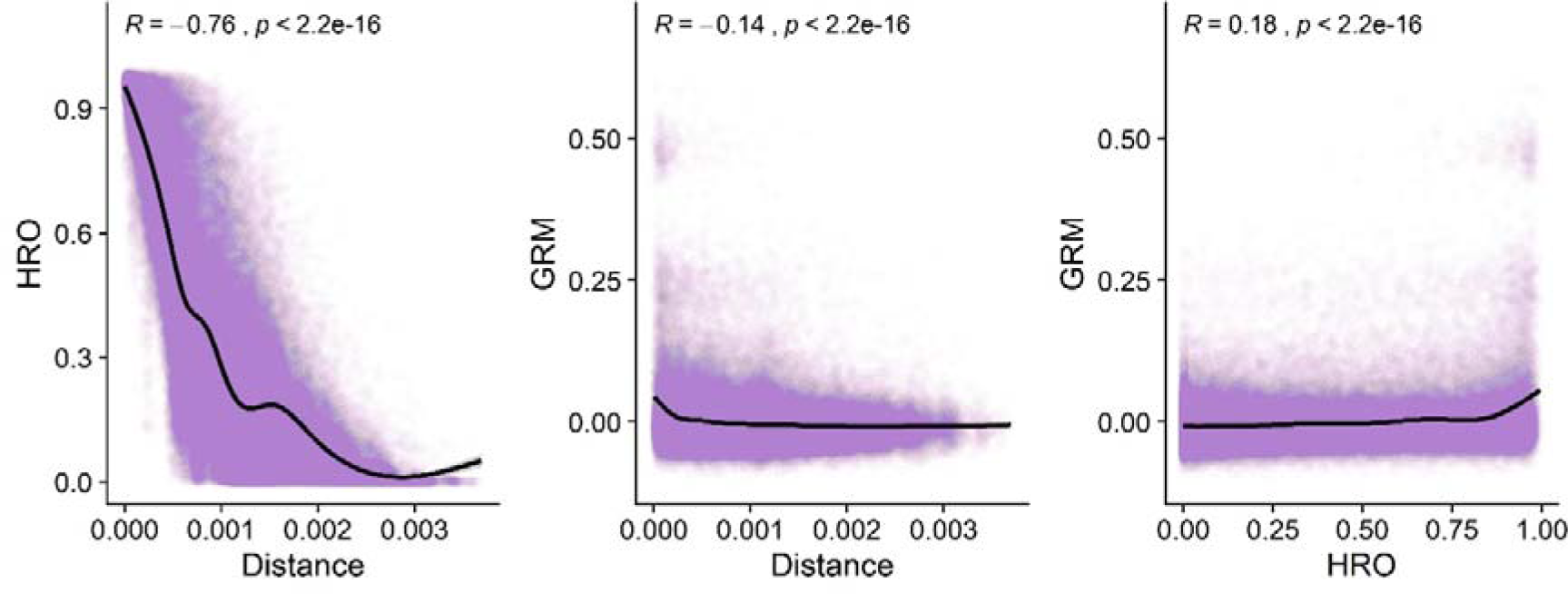
Correlation among pairwise values in lifetime point location distances, home range overlap matrix, and genomic relatedness matrix. The data have been fitted with a Generalised Additive Model (GAM) smooth.

**Supplementary Figure 2:**
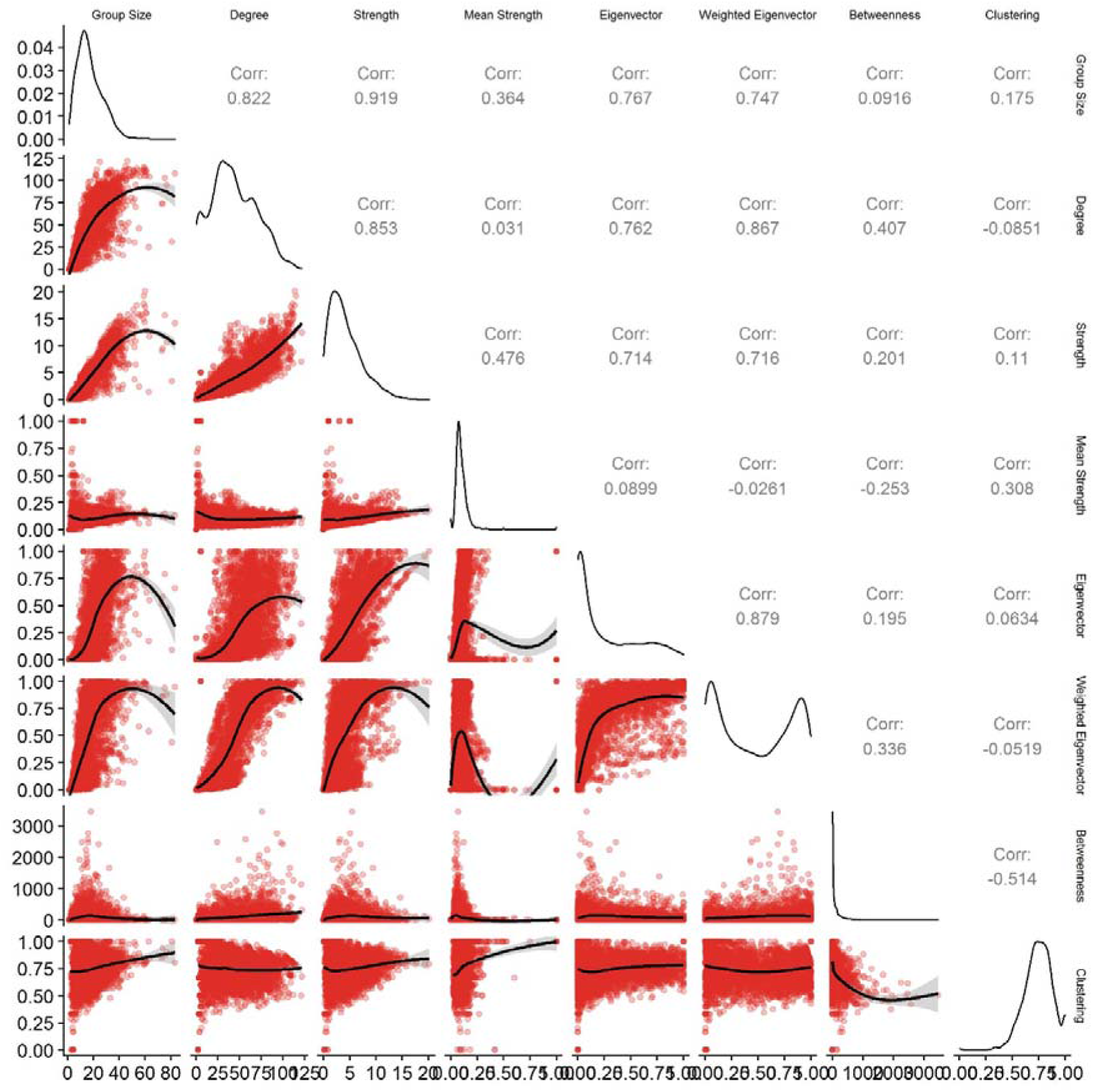
Pairwise correlations among network position response variables. Values were transformed, scaled to have a mean of 0 and a standard deviation of 1, with outliers removed, before analysis. The data have been fitted with a Generalised Additive Model (GAM) smooth.

**Supplementary Figure 3:**
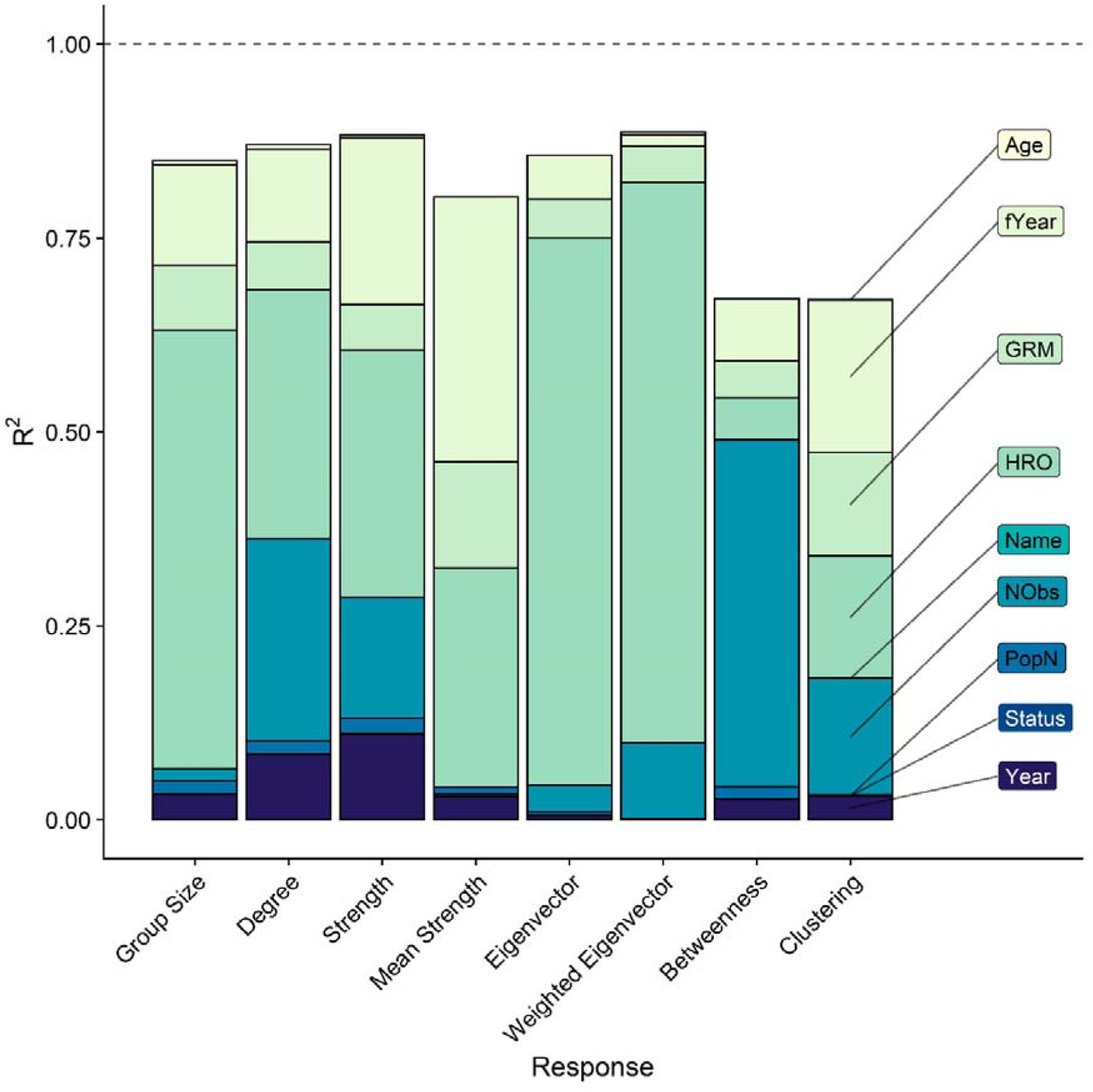
Variance accounted for by each variable for all eight network position measures, expressed as contribution to R^2^ in the models with home range overlap. This output differs from Figure 3B in the main text because the model did not include the SPDE random effect. Different shades correspond to different variables. fYear = year as a categorical random effect. GRM = Genomic Relatedness Matrix. HRO = home range overlap. Name = individual identity. NObs = number of observations (i.e., sampling bias). PopN = population size. Status = reproductive status. For all response variables, individual level variables (Age, Reproductive Status, Identity) had a negligible effect.

**Supplementary Figure 4:**
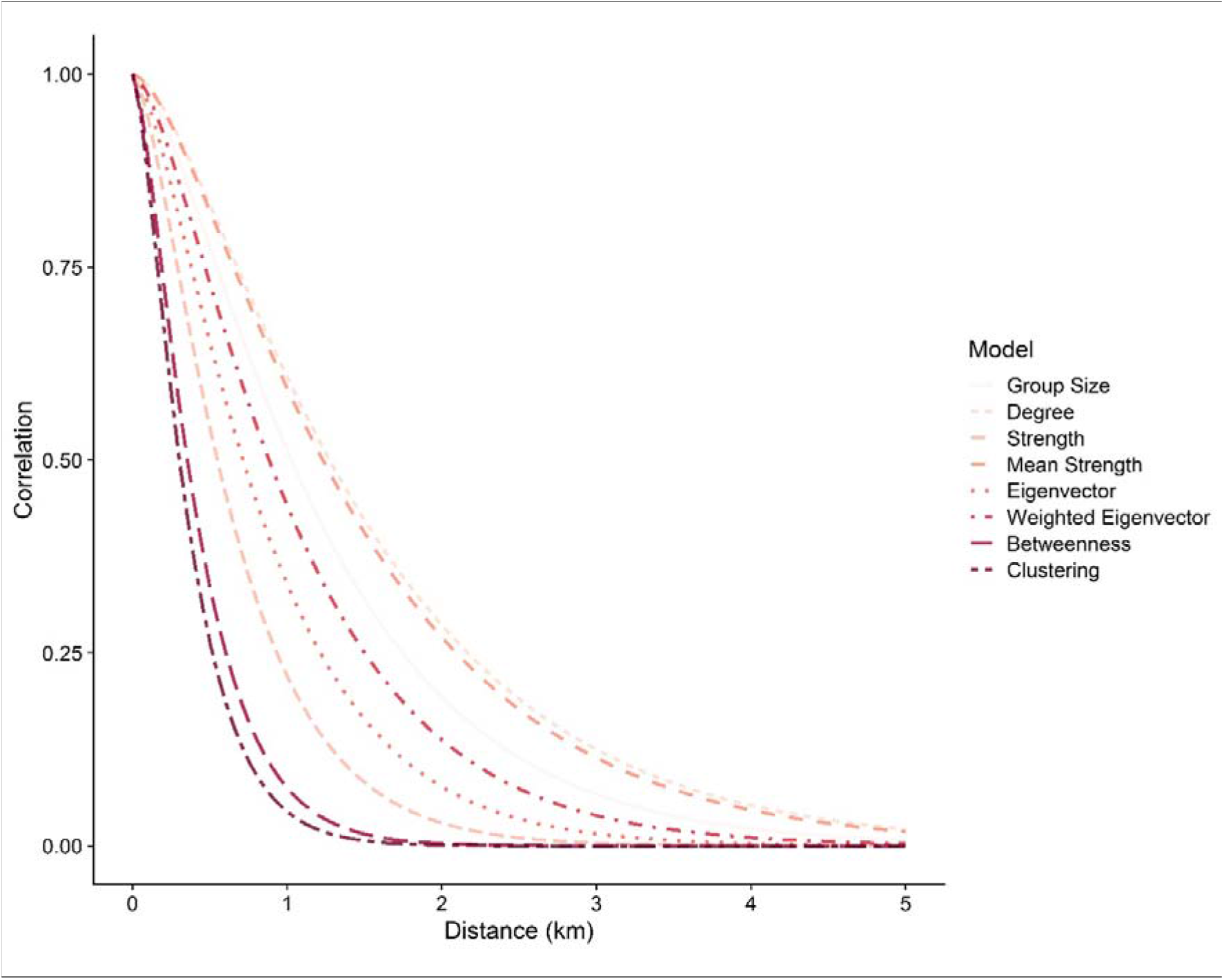
The INLA SPDE autocorrelation ranges for each response variable. Different shades and line types correspond to different response variables. The X axis is in kilometres.

**Supplementary Figure 5:**
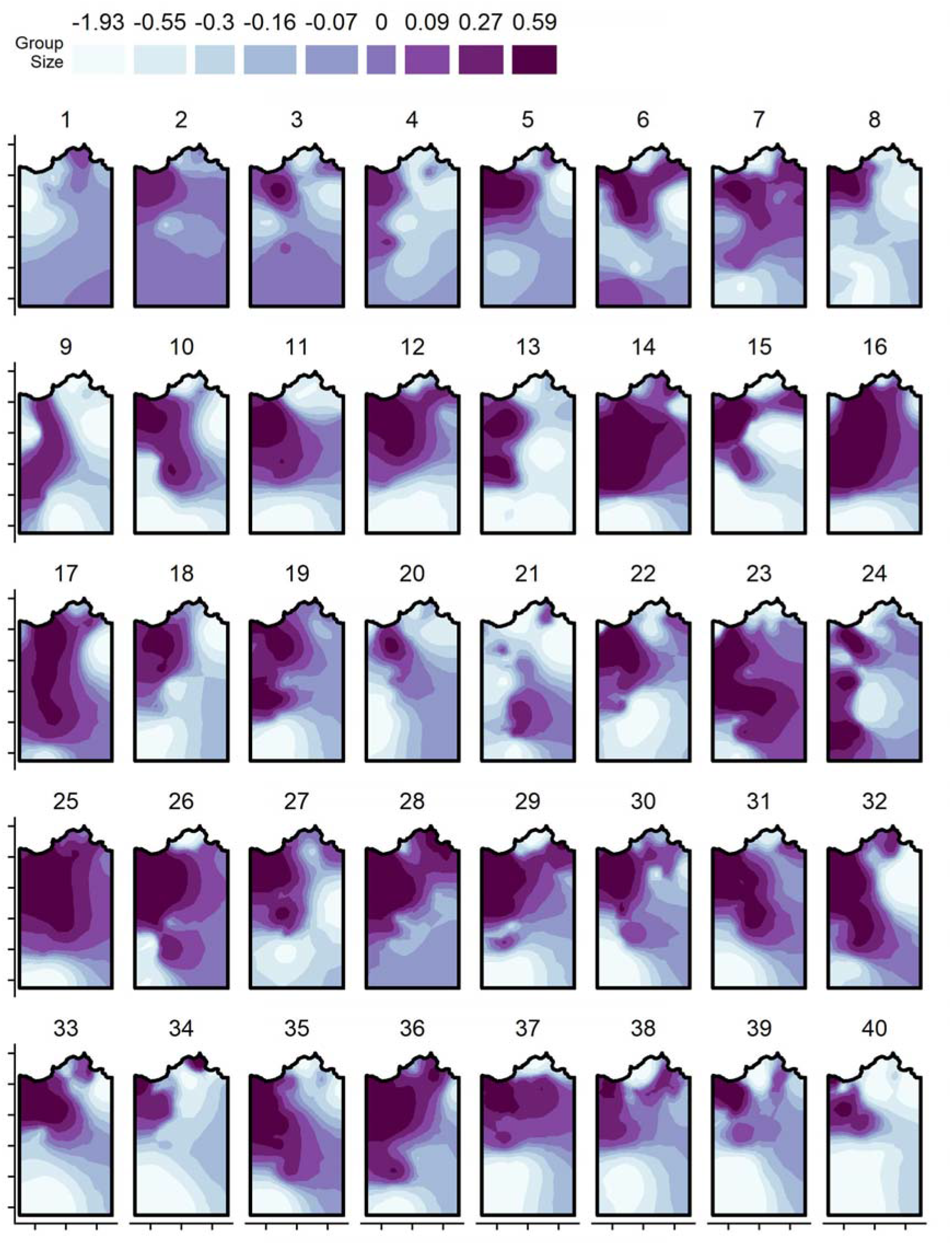
Annual spatial fields for the SPDE random effect for group size, taken from the INLA animal models and based on annual centroid point locations. Darker colours correspond to greater values. Each axis tick corresponds to 1km distance; for the values associated with the Easting and Northings, see Figure 1 in the main text.

**Supplementary Figure 6:**
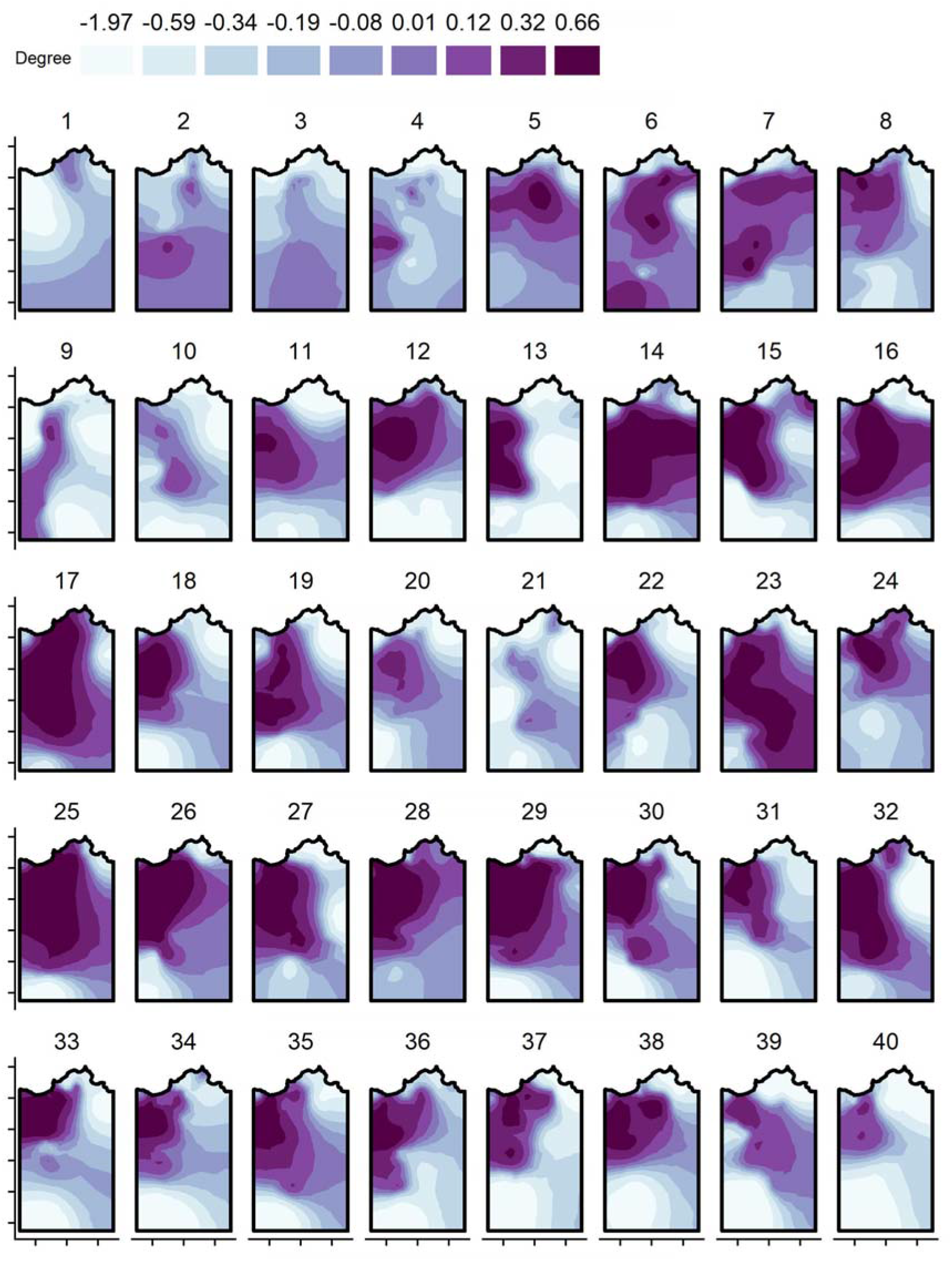
Annual spatial fields for the SPDE random effect for degree centrality, taken from the INLA animal models and based on annual centroid point locations. Darker colours correspond to greater values. Each axis tick corresponds to 1km distance; for the values associated with the Easting and Northings, see Figure 1 in the main text.

**Supplementary Figure 7:**
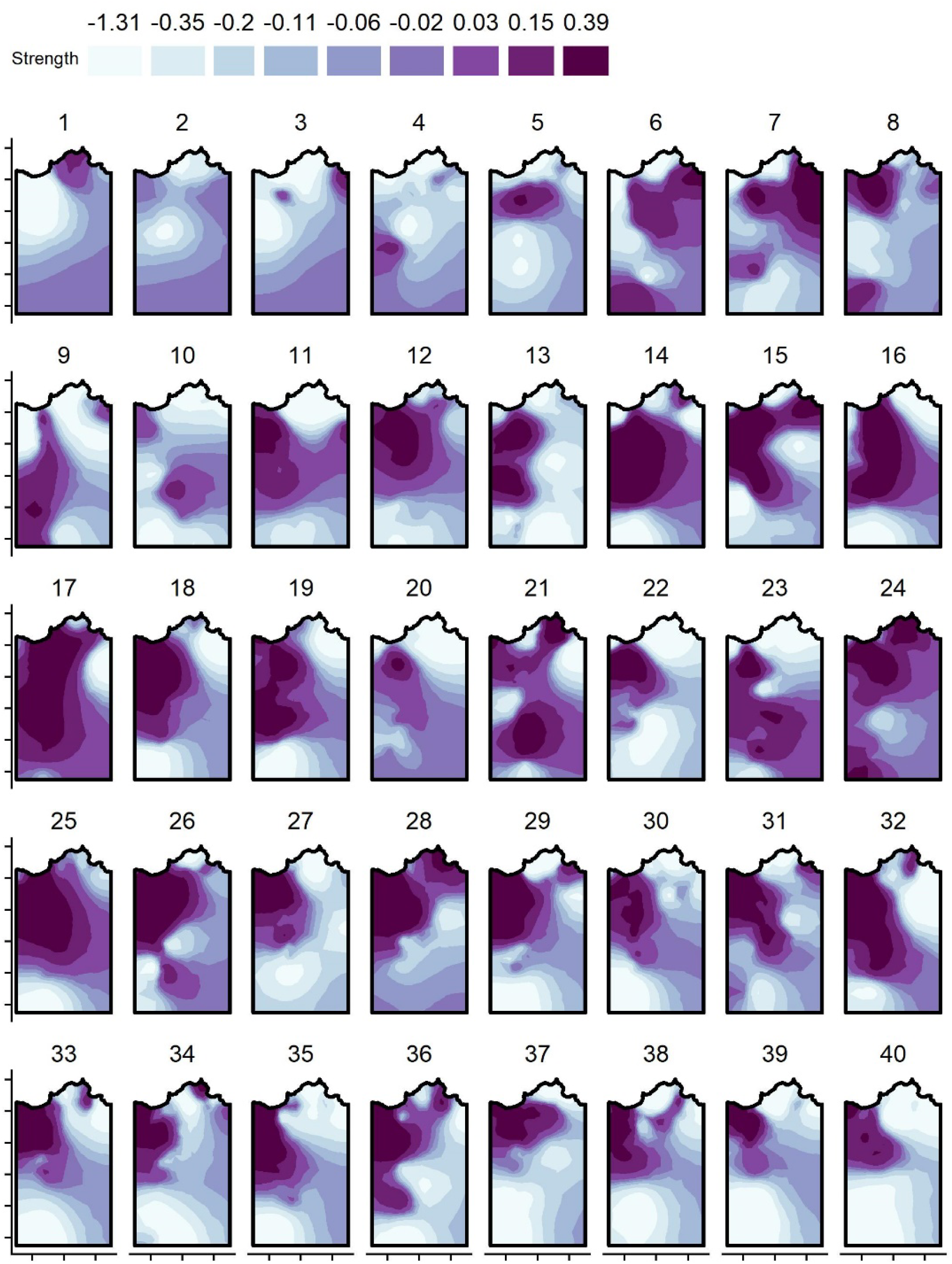
Annual spatial fields for the SPDE random effect for strength centrality, taken from the INLA animal models and based on annual centroid point locations. Darker colours correspond to greater values. Each axis tick corresponds to 1km distance; for the values associated with the Easting and Northings, see Figure 1 in the main text.

**Supplementary Figure 8:**
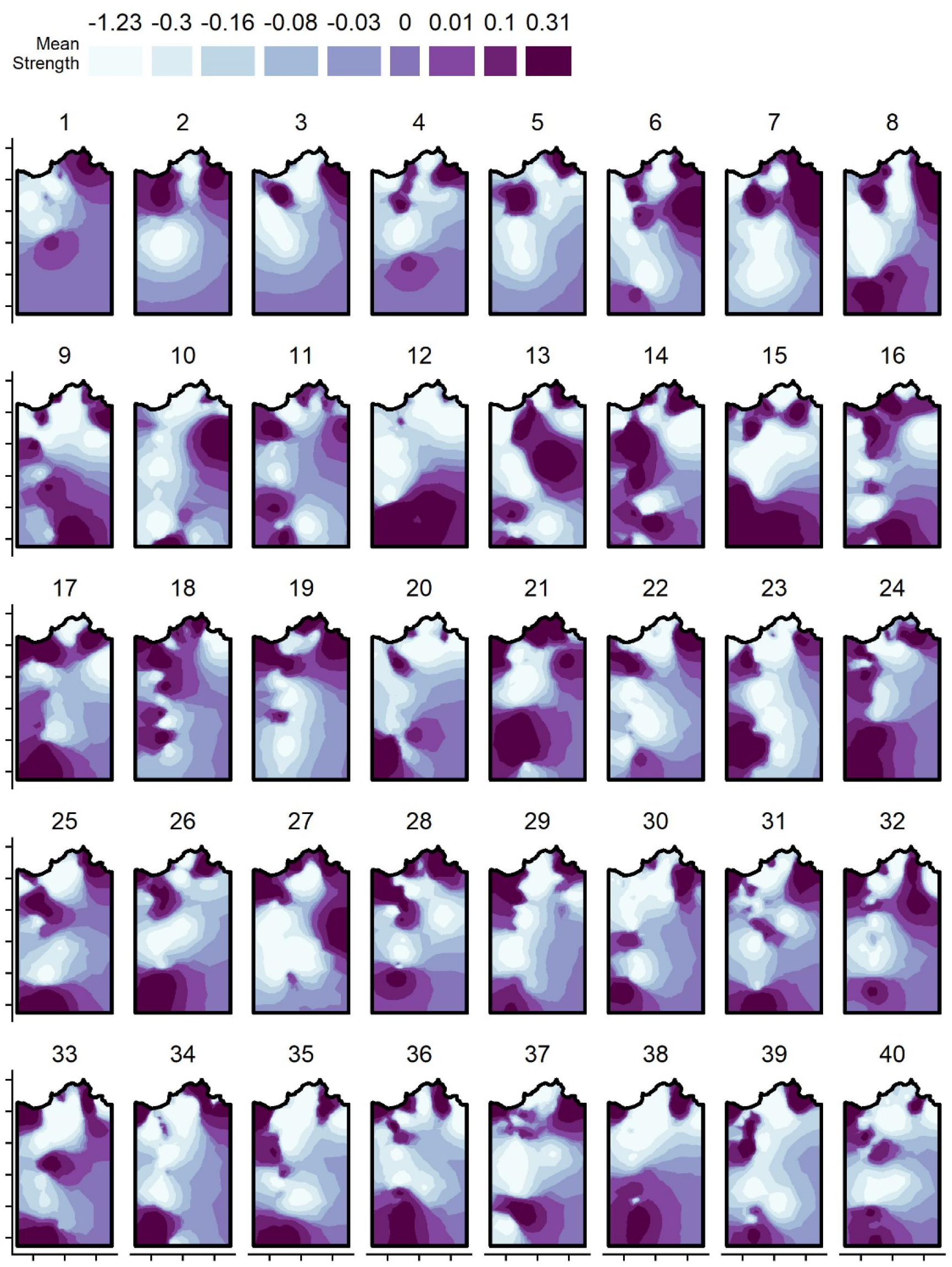
Annual spatial fields for the SPDE random effect for mean strength centrality, taken from the INLA animal models and based on annual centroid point locations. Darker colours correspond to greater values. Each axis tick corresponds to 1km distance; for the values associated with the Easting and Northings, see Figure 1 in the main text.

**Supplementary Figure 9:**
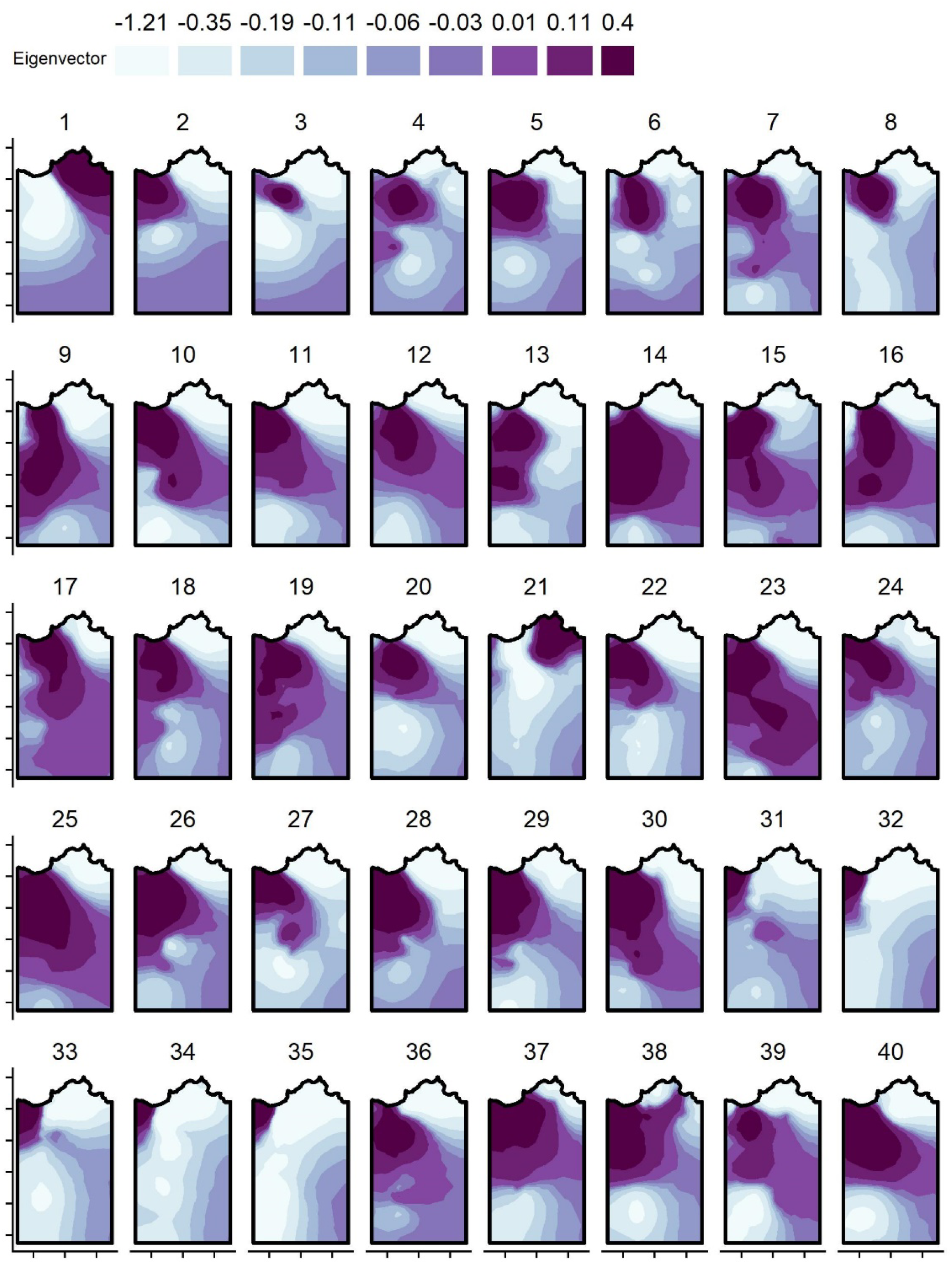
Annual spatial fields for the SPDE random effect for Eigenvector centrality, taken from the INLA animal models and based on annual centroid point locations. Darker colours correspond to greater values. Each axis tick corresponds to 1km distance; for the values associated with the Easting and Northings, see Figure 1 in the main text.

**Supplementary Figure 10:**
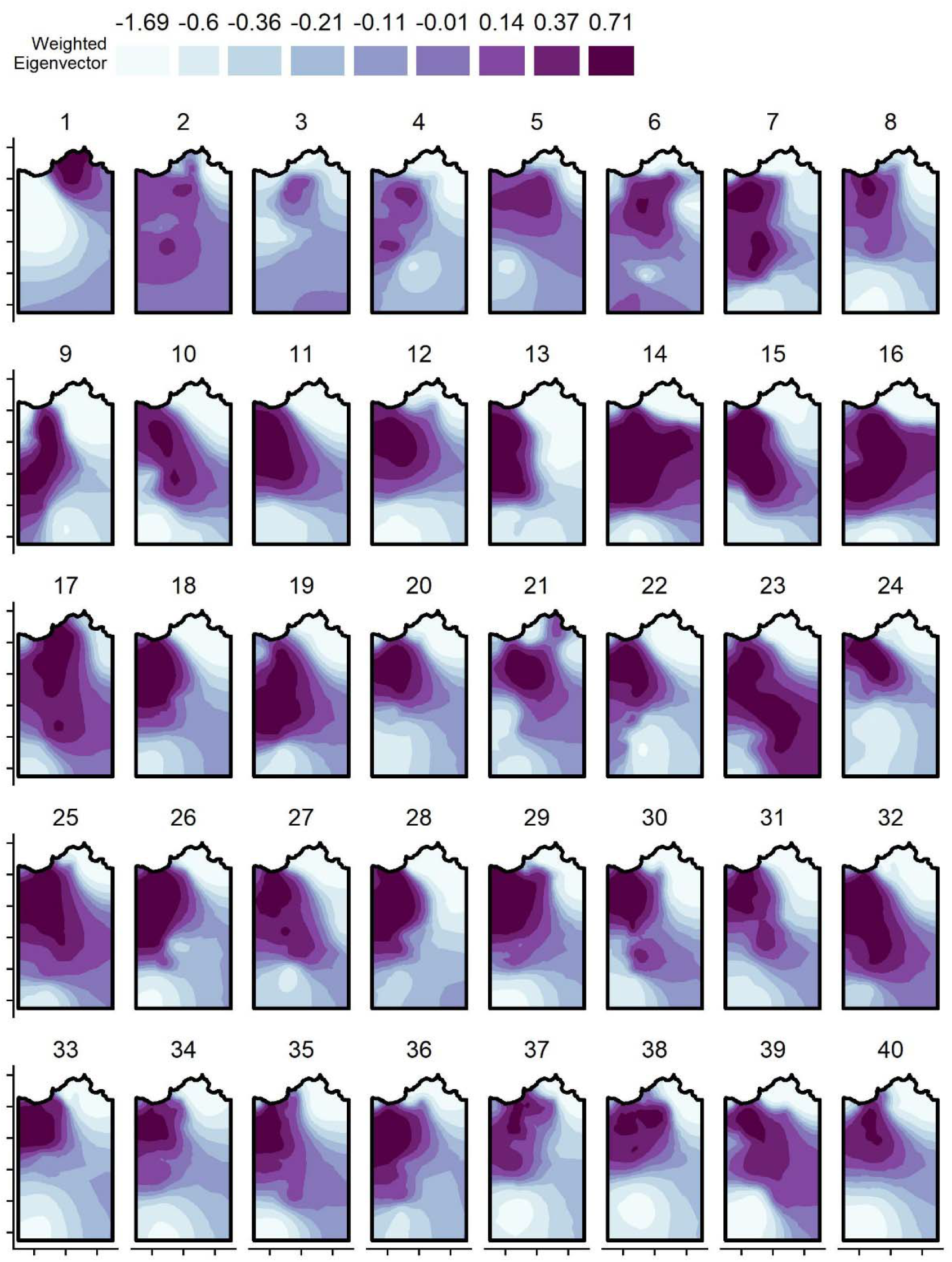
Annual spatial fields for the SPDE random effect for weighted Eigenvector centrality, taken from the INLA animal models and based on annual centroid point locations. Darker colours correspond to greater values. Each axis tick corresponds to 1km distance; for the values associated with the Easting and Northings, see Figure 1 in the main text.

**Supplementary Figure 11:**
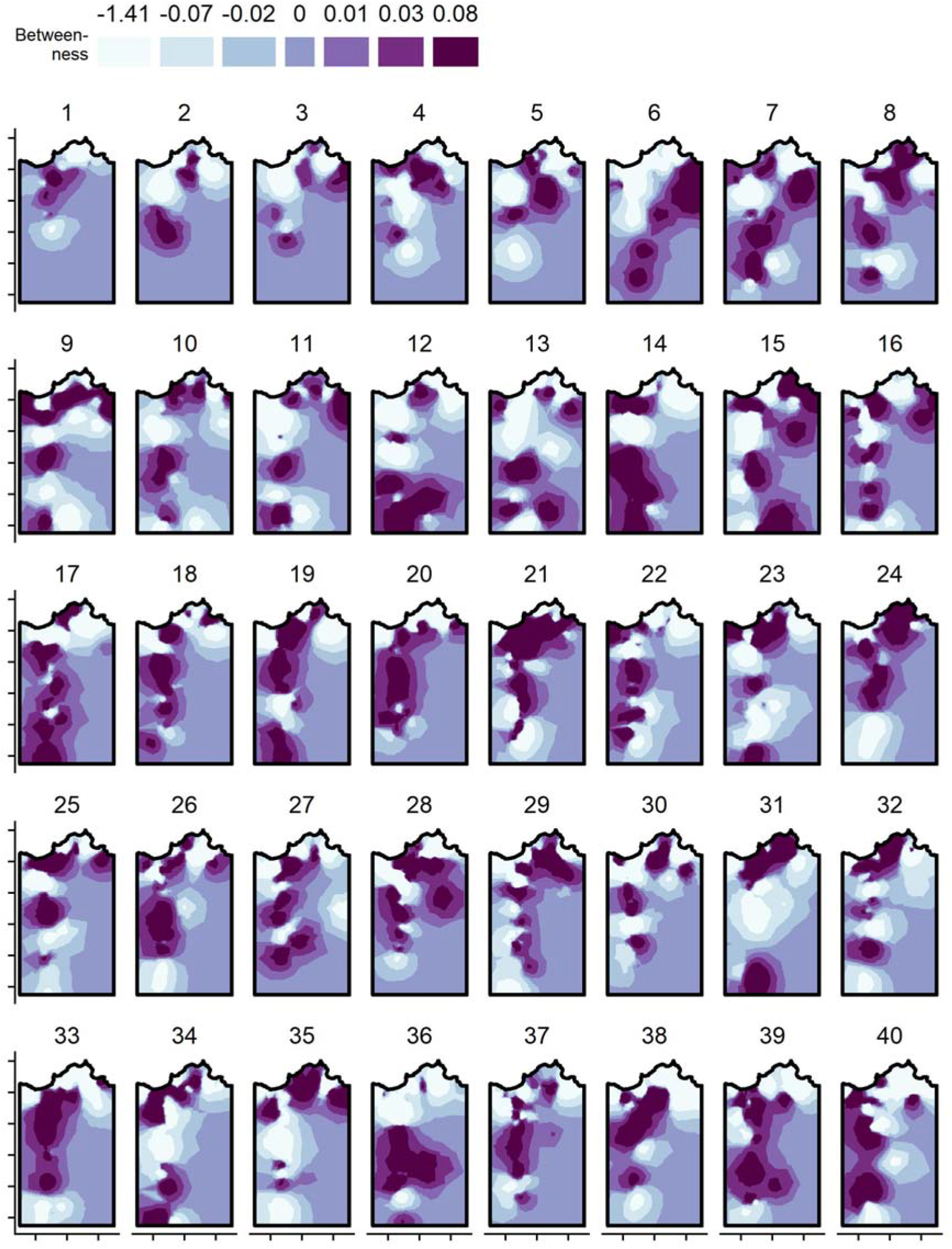
Annual spatial fields for the SPDE random effect for betweenness centrality, taken from the INLA animal models and based on annual centroid point locations. Darker colours correspond to greater values. Each axis tick corresponds to 1km distance; for the values associated with the Easting and Northings, see Figure 1 in the main text.

**Supplementary Figure 12:**
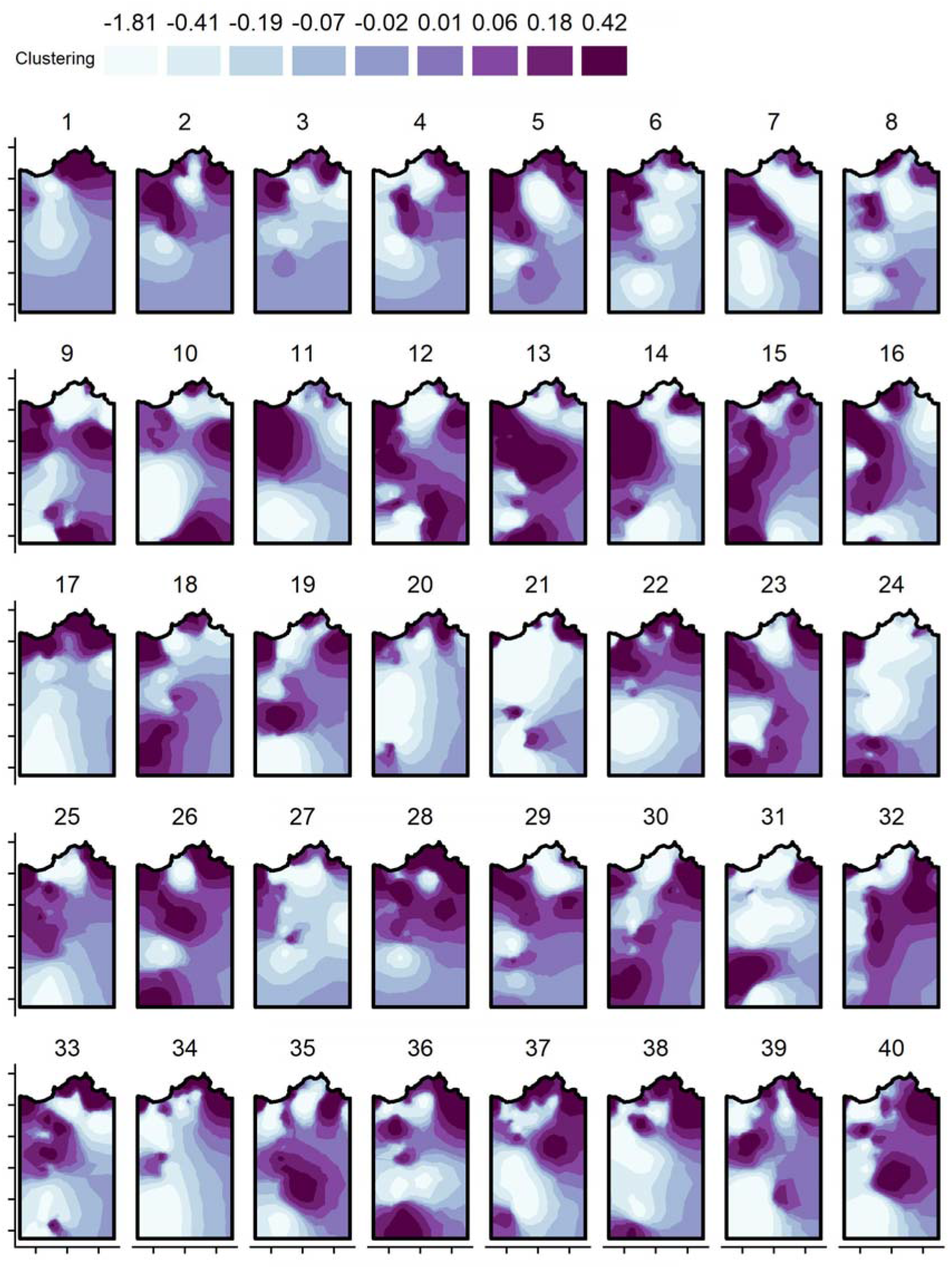
Annual spatial fields for the SPDE random effect for clustering, taken from the INLA animal models and based on annual centroid point locations. Darker colours correspond to greater values. Each axis tick corresponds to 1km distance; for the values associated with the Easting and Northings, see Figure 1 in the main text.

